# Coexisting but dissociable place and spatial view codes in the primate hippocampus

**DOI:** 10.64898/2026.07.20.739519

**Authors:** Seuk-Hwan Shin, Heung-Yeol Lim, Su-Min Lee, Jae-Min Seol, Bona Lee, Choong-Hee Lee, Yuji Naya, Joonyeol Lee, Inah Lee

**Author notes:** **Corresponding author:** Inah Lee, Joonyeol Lee.

## Abstract

In rodents, hippocampal activity is classically described in terms of position-anchored place cells, whereas nonhuman primate work has emphasized spatial view cells, in which activity depends on the region of space being viewed. This contrast has fueled a longstanding debate over whether primate and rodent spatial codes are biologically distinct. We recorded single units in the hippocampus of macaques as they navigated linear tracks in two visually distinct contexts, intermittently making context-dependent object choices. Using a generalized linear model to dissociate position coding from spatial view coding, we identified three coexisting populations in the macaque hippocampus: Position cells, View cells, and Conjunctive cells encoding both variables. Position cells were prevalent and showed rodent-like properties—localized firing, directional tuning, and contextual remapping—even after accounting for visual sampling, whereas View cells stably encoded allocentric visual space. Population decoding confirmed this dissociation at the ensemble level. Robust position codes thus coexist with, yet remain dissociable from, view codes in the primate hippocampus, suggesting that the apparent scarcity of place-like responses in earlier primate work reflects methodological and task-related differences rather than a fundamental species-level divergence.

## Introduction

The hippocampus is fundamentally involved in episodic memory and spatial navigation^1,2^. Rodent hippocampal place cells, which fire selectively as an animal traverses specific locations, led to the influential view that the hippocampus constructs an internal cognitive map of space. Building on this framework, subsequent work extended cognitive-map-based models of hippocampal function from rodents to nonhuman primates and humans^1,3–6^, shaping a broad research field centered on spatial coding and navigation. As this line of research has developed, studies in primates and humans^7–12^, often using virtual navigation and single-unit recordings or neuroimaging, have substantially advanced our understanding of how the hippocampus and its connected networks support spatial cognition across species.

Despite this widely adopted framework, the nature of spatial coding in primates remains unresolved. In rodents, hippocampal activity is typically described in terms of position-anchored place coding, whereas studies in nonhuman primates have often emphasized spatial view coding^11–18^, in which neurons fire according to the part of the environment being looked at rather than the animal’s physical location. Repeated reports of view-dominant coding, together with the importance of active visual sampling in primates with advanced foveal vision^19–24^, have led to the proposal that spatial representations in the primate hippocampus may be organized differently from those classically described in rodents. Interestingly, previous nonhuman primate studies have often failed to demonstrate place cells with the same robustness and spatial precision characterized in rodents, along with their defining features such as directional selectivity^25–28^ and contextual remapping^29–33^, across settings ranging from free navigation in open cages^15,16,34,35^ to linear tracks^36,37^ and virtual space^11,12,38^. These discrepancies have fueled an ongoing controversy: is the primate hippocampal code fundamentally distinct from the location-based representations found in rodents?^19,20,22,24^

Recent physiological evidence suggests that the primate hippocampus may instead be governed by mixed selectivity, in which individual neurons encode multiple variables concurrently rather than a single spatial quantity in isolation^11,15–17^. In this coding scheme, individual neurons encode multiple variables concurrently rather than a single spatial quantity in isolation. Studies employing multivariate models to account for diverse variables, such as location, spatial view, and facing direction have found that a larger proportion of hippocampal neurons exhibit mixed selectivity. Notably, only a small portion of hippocampal neurons has been reported as location-selective and are typically confounded with view direction or other behavioral variables, precluding the identification of pure position-dependent coding^11,15–17^. This prevalence of view-centric, multiplexed coding has reinforced the hypothesis that the primate hippocampal code is fundamentally distinct from the canonical spatial code observed in rodents^19–24^.

Despite these discrepancies, we hypothesized that the primate hippocampus can express robust, rodent-like spatial representations when specific experimental and analytical constraints are met. We proposed that such representations emerge under a combination of conditions: task demands necessitating active context sampling, exposure to a visually enriched virtual environment, and an analytical approach capable of separating position from the spatial view signal with which it covaries during navigation. To evaluate this hypothesis, we performed single-unit recordings in the hippocampus of macaque monkeys as they navigated a virtual linear track in two visually distinct contexts, with intermittent context-dependent object choices. To separate position coding from spatial view coding, we applied a generalized linear model at the single-neuron level, together with a field-of-view-based reconstruction of three-dimensional gaze. Our analysis revealed a heterogeneous neural population that includes both conjunctive cells and “canonical” cells, the latter exhibiting selective tuning for either position or spatial view that was independent of confounding variables. Collectively, these results suggest that the long-debated differences between rodent and primate hippocampal codes may reflect methodological and task-related factors at least as much as fundamental biological differences, helping to bridge a decades-old gap in comparative neuroscience.

## Results

### A VR linear-track navigation paradigm and field-of-view-based 3D gaze reconstruction

Two head-fixed monkeys (N and Y) navigated a virtual linear track using a joystick while hippocampal activity was recorded (Fig. 1A, Supplementary Fig. 1). On each lap, one of two visually distinct contexts, Forest or City, was presented pseudorandomly (Fig. 1B), and the animals shuttled along the track between outbound and inbound runs (Fig. 1C). At four predefined locations on the track, locomotion was halted and the monkeys were required to choose the object associated with the current context to obtain a water reward (Supplementary Fig. 2A). After training, both animals performed this context-dependent object-choice task at a high level across sessions and contexts (Supplementary Fig. 2B).

**Figure 1.**
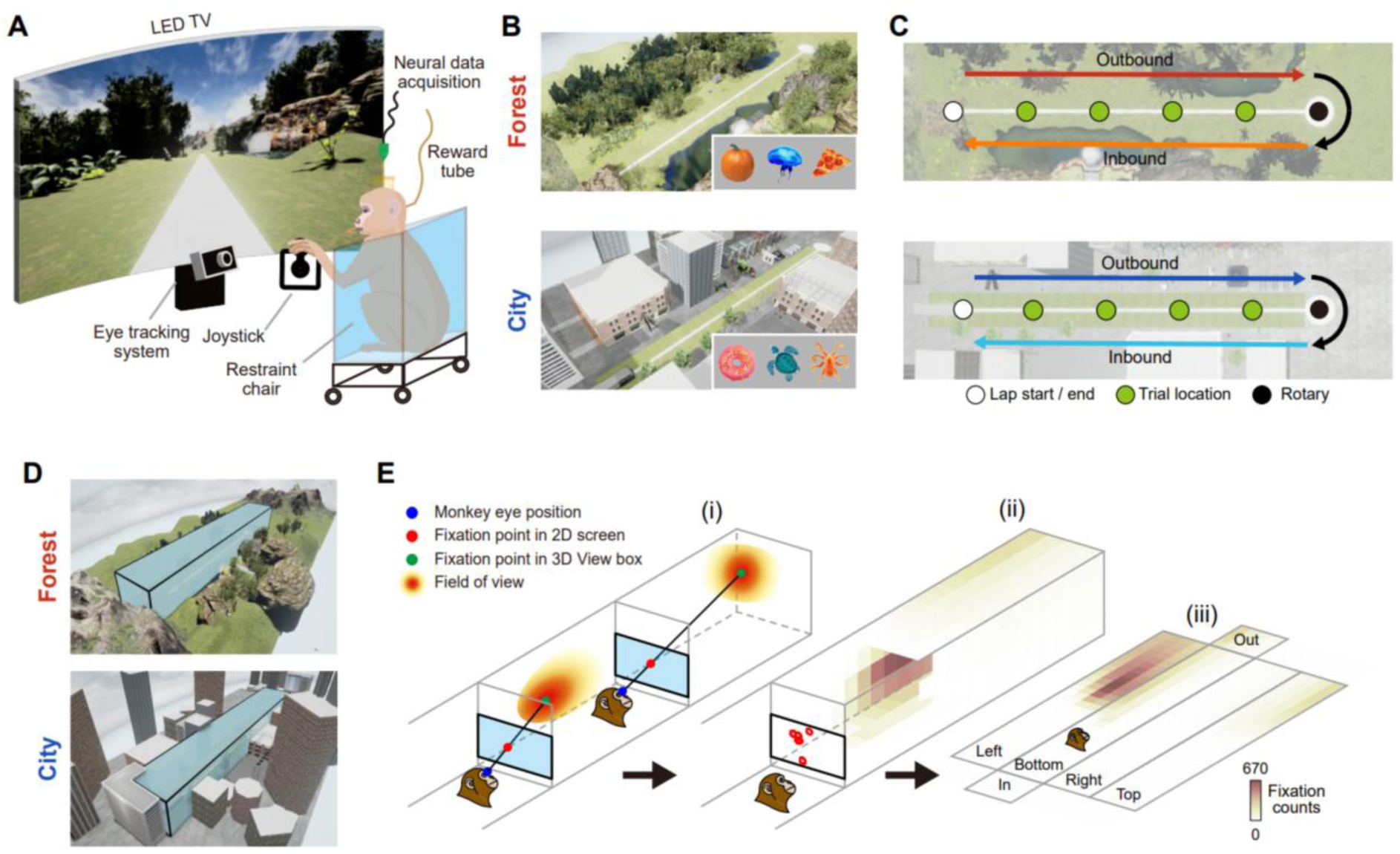
Virtual linear-track navigation task and 3D gaze reconstruction. **(A)** Schematic illustration of the virtual reality setup for the behavioral task and neural recording. Head-fixed monkeys manipulated a joystick to navigate along a linear track in virtual contexts presented on a LED screen. Eye movements were recorded simultaneously with neural spiking activity. **(B)** Overview of the Forest (top) and City (bottom) contexts, illustrating their landmark configurations. Insets show the three distinct objects uniquely associated with each context. **(C)** Bird’s-eye view of the linear tracks in the two contexts. Arrows indicate outbound and inbound journeys. Green circles denote trial locations where the monkeys were stopped for context-dependent object choices. Monkeys experienced a 180° rotation at the rotary (black circle) to start inbound journeys. **(D)** Schematic of the virtual “view box” encompassing the linear track, used for 3D gaze reconstruction. **(E)** Methodology for mapping 2D gaze to 3D space. (i) Fixation points on the 2D screen were projected onto the virtual view box to determine 3D fixation points. (ii) Aggregated 3D fixation counts were mapped onto the surfaces of the view box, and then (iii) the view box was unfolded into a flat 2D gaze map.

Because primate spatial coding has often been interpreted through the lens of visual sampling, we reconstructed gaze behavior in three-dimensional space using a field-of-view framework applied to non-saccadic fixations in the eye-movement tracking data (Supplementary Fig. 3). We first defined a virtual view box enclosing the environment (Fig. 1E), and then each non-saccadic fixation was expanded into a Gaussian-weighted field of view and projected onto the surfaces of the virtual view box (Fig. 1F). This procedure generated continuous gaze maps that captured which regions of the environment were sampled visually during navigation. Gaze sampling patterns differed reliably between Forest and City contexts, and context identity was accurately decoded from the gaze maps for both monkeys (Supplementary Fig. 4), indicating that visual sampling was structured in a context-dependent manner.

### Position and view codes coexist but are dissociable in the primate hippocampus

To determine whether hippocampal neurons encoded position, spatial view, or both, Poisson generalized linear models (GLMs) were fit separately in each of the four task conditions defined by context and travel direction: Forest-Outbound (F-O), Forest-Inbound (F-I), City-Outbound (C-O), and City-Inbound (C-I) (Fig. 2A). For each neuron, position, view, and conjunctive models were compared by cross-validated encoding performance, and neurons were classified according to the model that best explained their activity. Representative examples showed that each class was well captured by its corresponding model, with reconstructed model rate maps closely matching the actual firing patterns (Fig. 2B-D). Across all recorded single units (n = 113), 87.7% were classified as spatially modulated in at least one task condition, whereas the remainder were non-spatial (Null) in all conditions (Fig. 2E). All three classes were present in every condition (Fig. 2F), and their relative proportions did not differ significantly across conditions (χ²_(9)_ = 9.13, p = 0.43, chi-square test), indicating that the three coding schemes were consistently expressed across contexts and directions.

**Figure 2.**
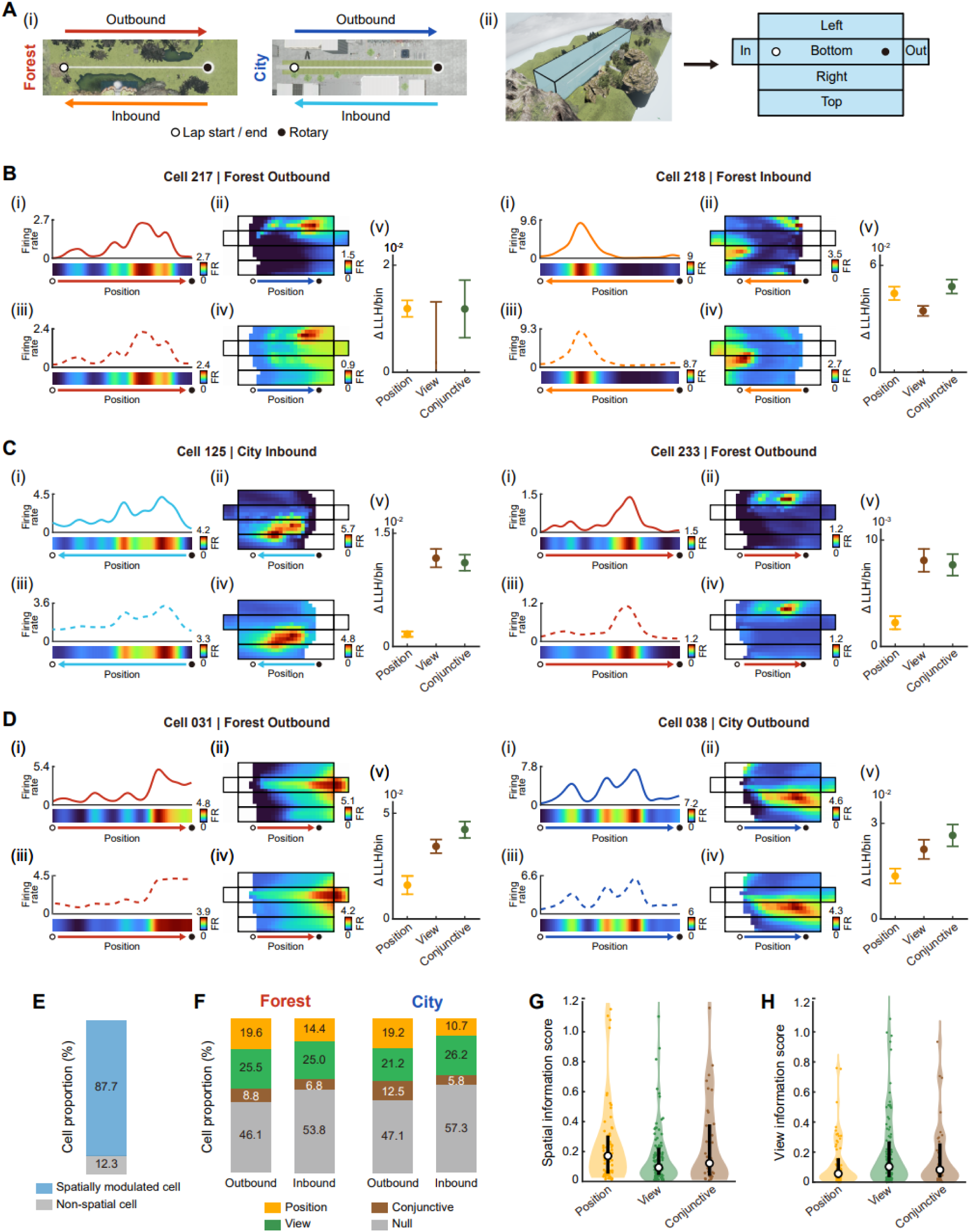
A generalized linear model dissociates Position, View, and Conjunctive cells in the primate hippocampus. **(A)** Schematic of the spatial coordinate frames used to construct position and view rate maps for the generalized linear model (GLM) analysis. (i) The four task conditions, defined by context (Forest, City) × direction (outbound, rightward; inbound, leftward). Task conditions are color-coded consistently throughout the figures as Forest-Outbound (F-O), red; Forest-Inbound (F-I), orange; City-Outbound (C-O), blue; City-Inbound (C-I), cyan. (ii) The virtual “view box” enclosing the linear track — used to reconstruct 3D gaze — unfolded into a flat layout. Its four side faces (Left, Bottom, Right, Top) and two end walls (In, Out) define the coordinate frame of the 2D view rate maps shown throughout the figures. **(B–D)** Representative neurons classified by the GLMs as Position cells (B), View cells (C), and Conjunctive cells encoding both position and view (D). For each cell, the top row shows empirical rate maps: firing rate as a function of position along the track (i) and across the unfolded view box (ii). The bottom row shows the corresponding modeled rate maps for position (iii) and view (iv). At right (v), encoding performance (ΔLLH/bin) for the position, view, and conjunctive models is shown (points, mean; error bars, S.E.M. across cross-validation folds); the best-fitting model defines each cell’s class. **(E)** Proportion of spatially modulated neurons — those classified as Position, View, or Conjunctive cells in at least one task condition across all recorded single units (n = 113). **(F)** Class composition of spatially modulated cells in each task condition, expressed as the percentage of cells classified as Position, View, Conjunctive, or Null. **(G, H)** Distributions of Skaggs spatial information scores (G) and view information scores (H) across GLM classes, shown as violin plots with overlaid jittered single-cell values (small filled circles), class medians (large open circles), and 95% bootstrap confidence intervals (error bars). One Position cell with an SI value > 1.2 was excluded from the SI distribution for visualization purposes.

To relate the model-based classes to more conventional spatial metrics, Skaggs’ spatial information^39^ (SI) and view information (VI) were compared across groups. The median SI was highest in Position cells (Position: 0.171; View: 0.093; Conjunctive: 0.122; Fig. 2G), whereas the median VI was highest in View cells (Position: 0.057; View: 0.103; Conjunctive: 0.082; Fig. 2H), with Conjunctive cells exhibiting intermediate values on both measures. However, neither comparison reached statistical significance (SI: χ²_(2)_ = 4.63, p = 0.10; VI: χ²_(2)_ = 3.56, p = 0.17; Kruskal–Wallis test), and the distributions overlapped substantially across classes. These results indicate that marginal information scores alone are insufficient to cleanly separate the underlying coding types. They underscore the need for the model-based dissociation of position and spatial view coding.

### Position cells show stable and localized spatial tuning

We next characterized the firing properties of Position cells identified by the GLMs. Cells with high SI (> 0.15) showed sharply localized tuning, with spikes concentrated at circumscribed locations along the track across laps (Fig. 3A). Others with lower SI exhibited broader or multi-peaked tuning (Fig. 3B). Importantly, these cells were still classified by the GLMs as Position cells, indicating that low values of a marginal information metric did not preclude robust position-dependent coding.

**Figure 3.**
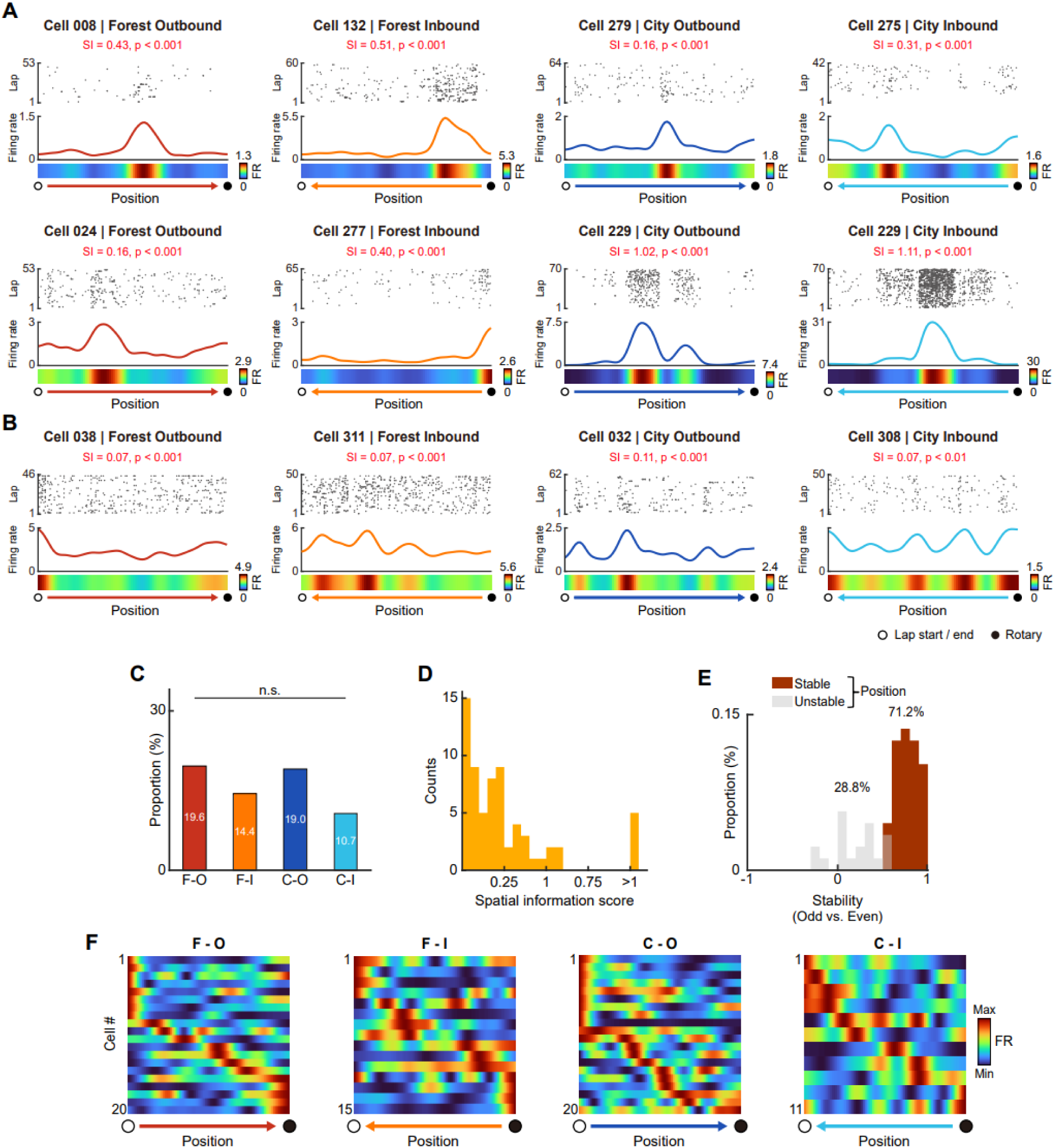
Position cells in the primate hippocampus during virtual navigation. **(A)** Representative Position cells with high spatial information (SI > 0.15). Two example cells were shown for each task condition. For each cell, three panels are displayed: a spike raster across laps as a function of track position (top), an averaged firing-rate map (FR; middle), and the corresponding firing-rate heat map (bottom). The arrow at the bottom indicates the direction of travel and is color-coded by task condition. Text in red denotes the SI score and its p-value for the displayed condition (permutation test against shuffled spike trains). **(B)** Representative Position cells with low spatial information (SI ≤ 0.15), showing weaker, multi-peaked position tuning. **(C)** Proportion of Position cells within each task condition (chi-square test; n.s., not significant). **(D)** Distribution of SI scores for Position cells (pooled across task conditions; rightmost bin, SI > 1). **(E)** Distribution of spatial firing stability for Position cells, quantified as Pearson correlation coefficients between position rate maps from odd and even laps. Cells with significantly stable firing relative to shuffled spike trains are shown in brown (permutation test, p < 0.05); unstable cells are shown in grey. **(F)** Population rate maps for Position cells in each task condition. Each row represents one cell’s peak-normalized position rate map.

The proportion of Position cells did not differ significantly across task conditions (χ²_(3)_ = 4.09, p = 0.25, chi-square test), corresponding to 11% to 20% of neurons per condition and 41.6% of all neurons in at least one condition (Fig. 3C). SI scores spanned a broad range (Fig. 3D), consistent with the coexistence of sharply tuned single-field responses and broader or multi-peaked tuning patterns within the same population. When we examined the tuning stability by correlating each cell’s rate maps between odd and even laps, 71.2% of Position cells showed significant stability (Fig. 3E), suggesting that position tuning was reliable within a session.

At the population level, Position-cell rate maps collectively tiled the linear track in every condition (Fig. 3F). Fields were overrepresented near the lap start/end region, corresponding to the point at which the animal entered a context at the beginning of the outbound run and left it at the end of the inbound run. Because context identity could change from lap to lap, this concentration near the context-entry region may reflect elevated demands on contextual identification upon entering each context. Taken together, these results indicate that the primate hippocampus contains a substantial population of neurons with stable and spatially structured position tuning during virtual navigation.

### Position codes undergo directional and contextual remapping

Position coding on a linear track can vary with direction of travel^25–28^ in rodents, so directional remapping was assessed by comparing each Position cell’s spatial firing across outbound and inbound runs within a given context (Fig. 4A). Among all cells coding position in at least one direction, 90.7% showed remapping across direction, whereas only 9.3% retained stable firing patterns (Fig. 4B). The majority of cells with direction-specific firing were classified as Position cells in only a single direction in both contexts, whereas 12.9% (Forest) and 14.8% (City) of the cells were classified as Position cells in both directions (Fig. 4C).

**Figure 4.**
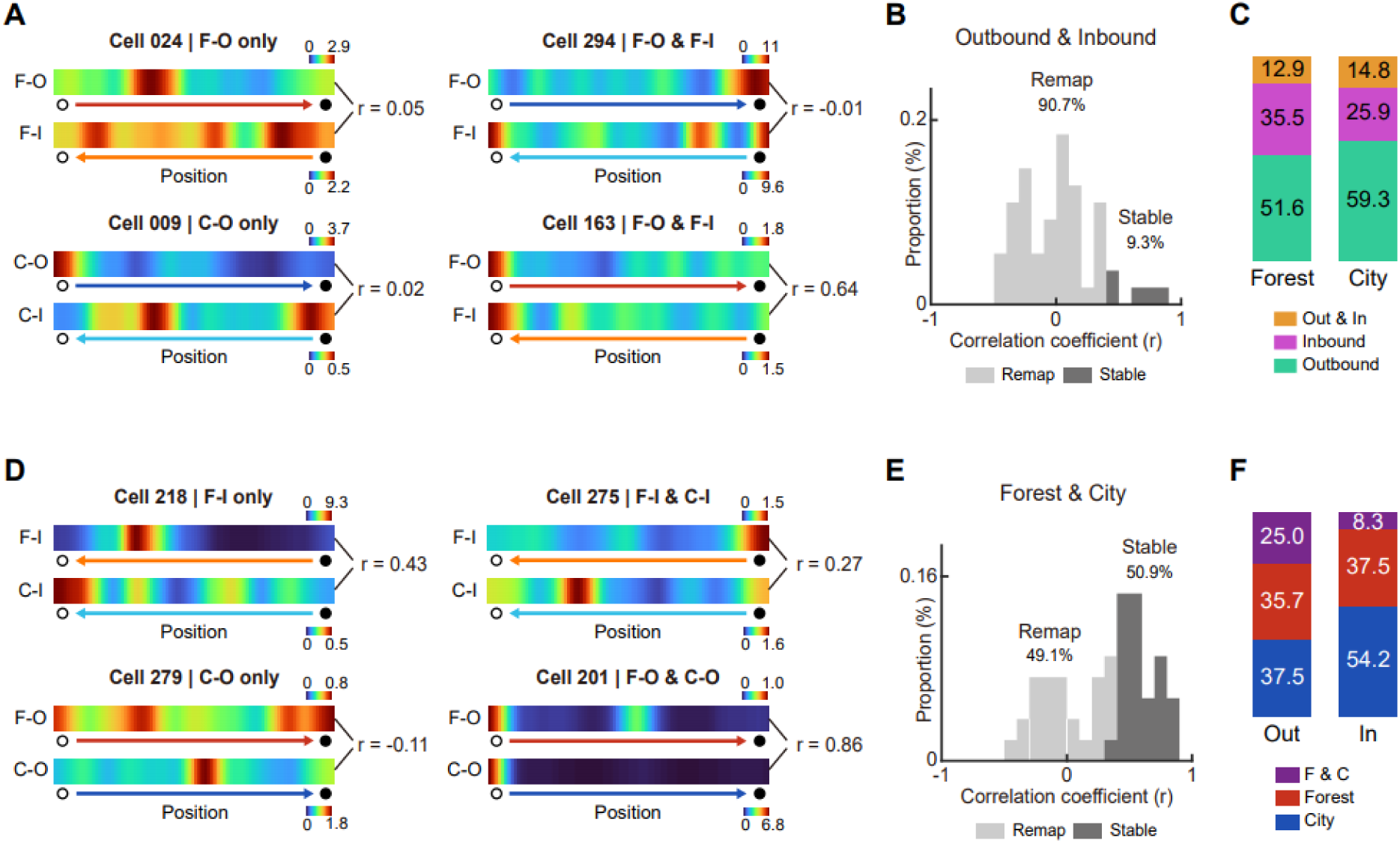
Directional and contextual remapping of position representations. **(A)** Example Position cells illustrating directional reorganization of position representations across travel directions within a context, except for the bottom-right cell whose firing pattern did not differ across directions. For each cell, position rate maps for the two opposite directions are shown together with the across-direction rate-map correlation (r). **(B)** Across-direction position rate-map correlations for all cells classified as Position cells in at least one of the two directions, pooled across within-context direction pairs (F-O with F-I, and C-O with C-I). Cells are classified as stable (dark gray) or remapping (light gray) based on a permutation test (p < 0.05). **(C)** Proportional composition of Position cells that code position in the outbound direction only, the inbound direction only, or in both directions (Out & In), shown separately for the Forest and City contexts. **(D)** Example Position cells illustrating contextual reorganization of firing patterns across contexts within the same direction, except for the bottom-right cell whose firing pattern did not differ across contexts. **(E)** Across-context position rate-map correlations for all cells classified as Position cells in at least one of the two contexts, pooled across within-direction pairs (F-O with C-O, and F-I with C-I). Cells are classified as stable (dark gray) or remapping (light gray) based on a permutation test (p < 0.05). **(F)** Proportional composition of Position cells that showed position coding in the Forest context only, the City context only, or in both contexts (F & C), shown separately for the outbound and inbound directions.

We next asked whether Position cells also remap between contexts by comparing each cell’s firing between Forest and City contexts within the same direction (Fig. 4D). Among cells coding position in at least one context, 49.1% showed remapping whereas 50.9% maintained stable firing across contexts (Fig. 4E). Most of the cells with context-dependent remapping were classified as Position cells in a single context, with only 25.0% (outbound) and 8.3% (inbound) classified in both contexts (Fig. 4F). Collectively, the spatial representations in the primate hippocampus reorganized systematically as a function of navigational direction and environmental context, indicating that hippocampal position codes in this task were dynamic and condition-dependent.

### View cells form stable representations of allocentric visual space

We next characterized the View cells identified by the GLMs. View cells fired according to the region of the environment sampled by gaze, represented as a three-dimensional rate map on the unfolded surfaces of the view box (Fig. 5A). As with Position cells, View cells ranged from sharply tuned units with concentrated view fields (VI > 0.15; Fig. 5A) to cells with broader but still structured tuning (Fig. 5B).

**Figure 5.**
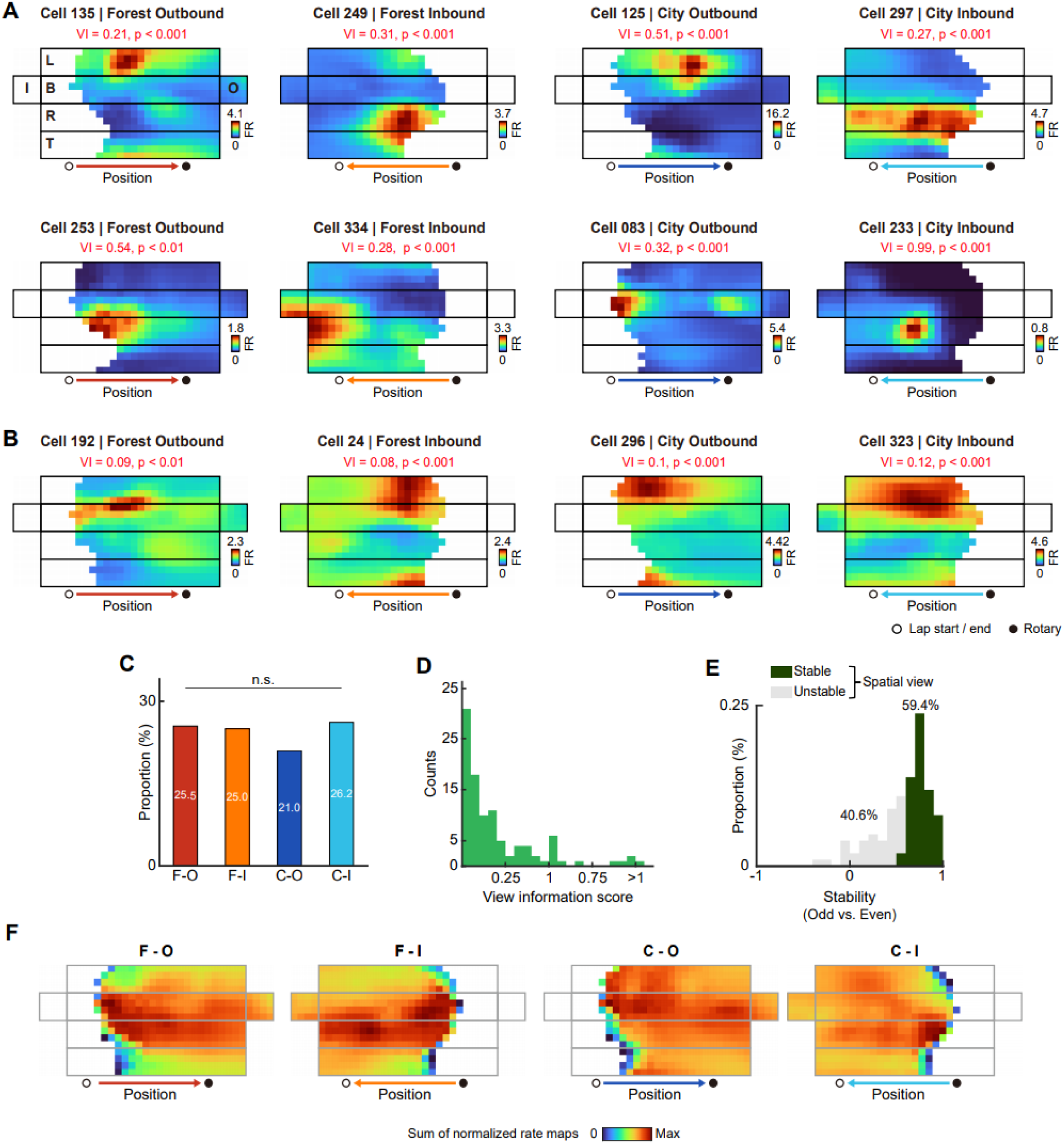
View cells in the primate hippocampus during virtual navigation. **(A)** Representative View cells with high view information (VI > 0.15). Two example cells are shown for each task condition. For each cell, the unfolded 2D view rate map is presented (box faces, end walls, arrow, and markers are as in Fig. 2A; color bar, firing rate). Text in red denotes the VI score and its permutation-test p-value for the displayed condition. **(B)** Representative View cells with low view information (VI ≤ 0.15), showing broader view tuning. **(C)** Proportion of View cells within each task condition (chi-square test; n.s., not significant). **(D)** Distribution of VI scores for View cells (pooled across task conditions; rightmost bin, VI > 1). **(E)** Distribution of view firing stability for View cells, quantified as Pearson correlation coefficients between view rate maps from odd and even laps. Cells classified as significantly stable relative to shuffled spike trains are shown in green (permutation test, p < 0.05); unstable cells are shown in grey. **(F)** Population view maps obtained by summing peak-normalized view rate maps across cells in each task condition.

The proportion of View cells likewise did not differ across conditions (χ²_(3)_ = 0.95, p = 0.81, chi-square test) and ranged from 21% to 26% per condition, corresponding to 50.4% of all neurons in at least one condition (Fig. 5C). VI scores were broadly distributed (Fig. 5D), and 59.4% of View cells showed significant odd–even lap stability (Fig. 5E). Population view maps (summing all View cells’ peak-normalized rate maps; Fig. 5F) broadly covered the unfolded view box in every condition, indicating distributed sampling of allocentric visual space rather than clustering around a small subset of locations. Together, these results identify a robust visuo-spatial code in the primate hippocampus. This code was reproducible across laps, broadly distributed across the environment, and dissociable from position coding at the single-unit level.

### View representations also reorganize across contexts

We then asked whether View cells are also modulated by environmental context by comparing each cell’s view-related firing between Forest and City within the same direction (Fig. 6A). Across all cells coding view in at least one context, 65.8% showed contextual remapping and 34.2% showed stable firing across contexts (Fig. 6B), which indicates that view representations reorganized across contexts more readily than position representations in the same comparison (cf. 49.1% remapping; Fig. 4F). About three-quarters of the context-modulated cells were classified as View cells in only one context, while the remainder were classified as View cells in both contexts (Fig. 6C).

**Figure 6.**
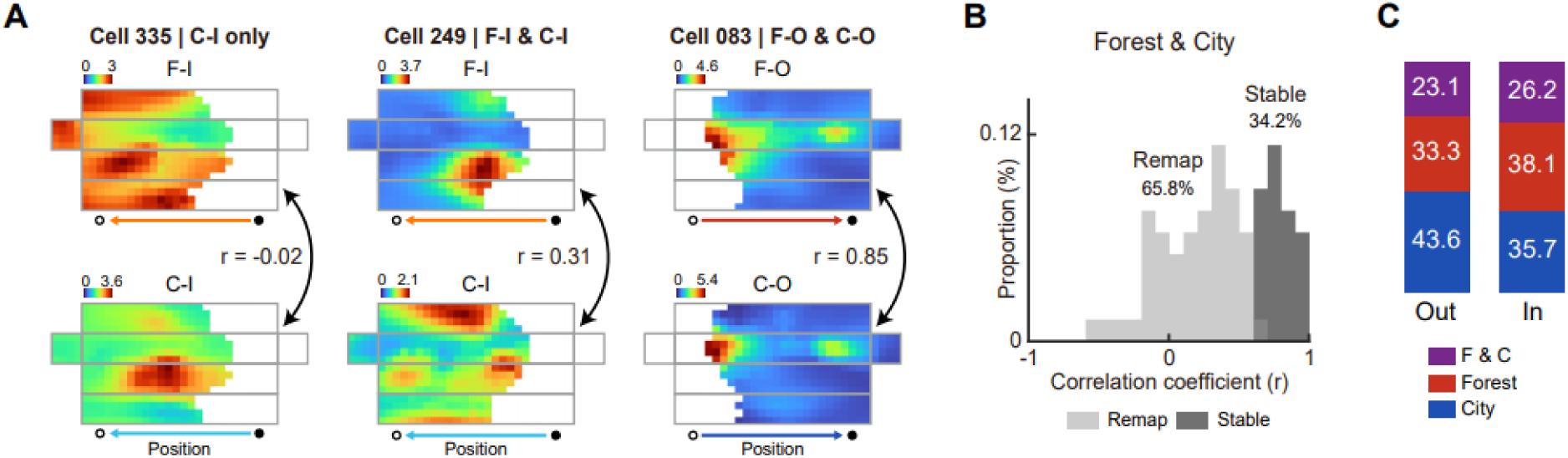
Contextual remapping of view representations. **(A)** Example View cells showing reorganized firing across contexts within a direction, while the rightmost cell showed stable view representations between contexts. For each cell, the unfolded view rate maps for each context (Forest, top; City, bottom) are shown with the across-context rate-map correlation coefficient (*r*). **(B)** Across-context view rate-map correlations for all cells that encode view in at least one context, pooled across within-direction pairs (F-O with C-O, and F-I with C-I). Cells are classified as stable (dark gray) or remapping (light gray) by a permutation test (p < 0.05). **(C)** Proportional composition of View cells that were classified as view coding in the Forest context only, the City context only, or in both contexts (F & C), shown separately for the outbound and inbound directions.

Taken together, context-dependent remapping is not unique to the position code: like Position cells, primate View cells reorganize their fields between the visually distinct environments. The two virtual environments were therefore distinguished along both positional and visual axes; that is, each context was associated with a distinct configuration of neural activity spanning both where the animal was and which part of the environment it sampled visually.

### Conjunctive cells jointly encode position and view

A third class of neurons, the Conjunctive cells, was best captured by the full model combining position and spatial view information and exhibited localized position fields together with structured view fields within the same neuron (Fig 7A). Example cells illustrated that position- and view-related tuning could coexist clearly within a single firing pattern. Conjunctive cells were the least frequent of the three spatial classes, representing 6%-12% of neurons per condition and 24.8% of all neurons in at least one condition (Fig. 7B). Their prevalence did not differ significantly across task conditions (χ²_(3)_ = 3.43, p = 0.33, chi-square test). Both SI and VI scores spanned broad ranges, and the corresponding distributions resembled those observed in Position and View populations (Fig. 7C; cf. Figs. 3D, 5D).

**Figure 7.**
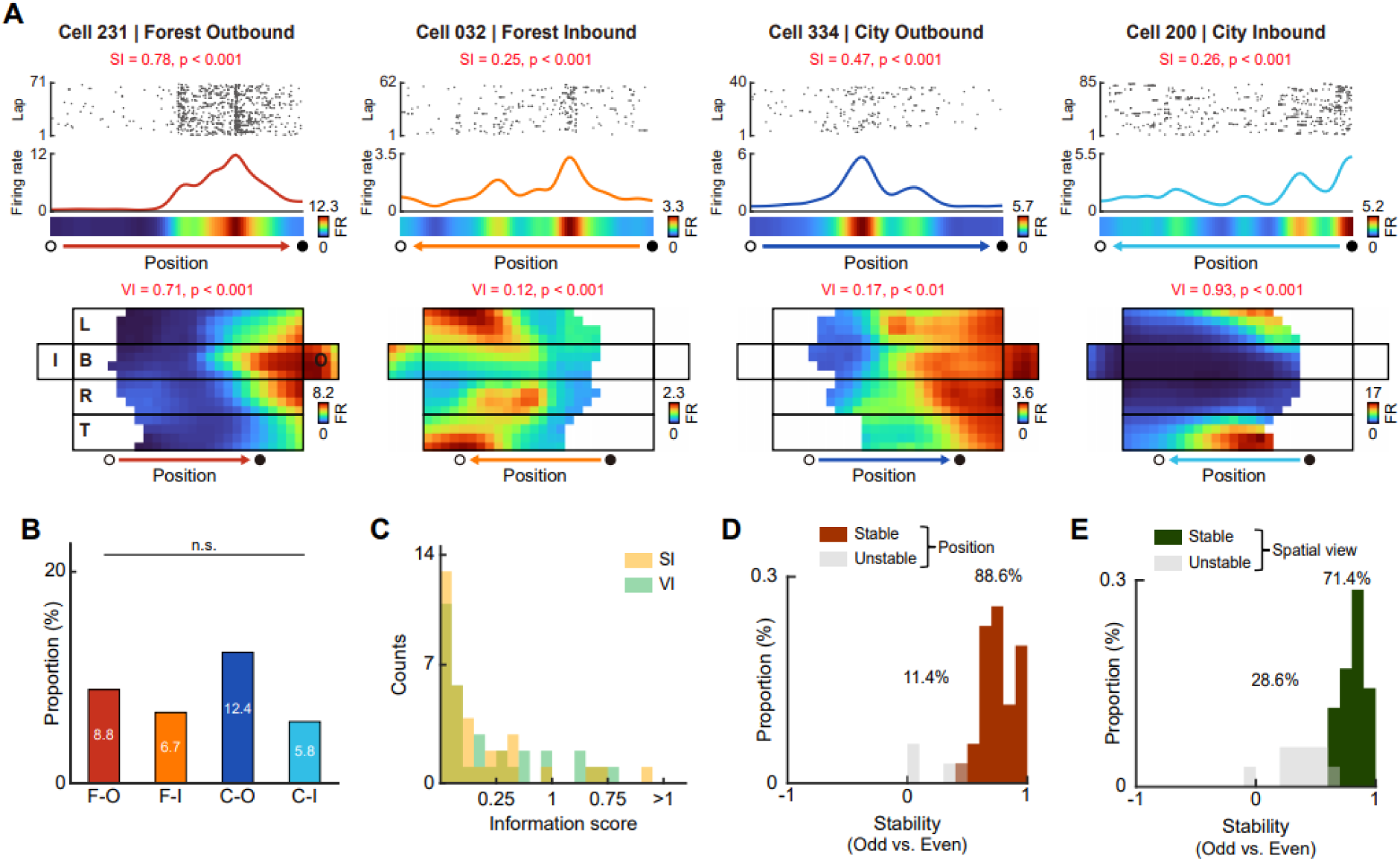
Conjunctive cells encoding both position and view information in the primate hippocampus during virtual navigation. **(A)** Representative Conjunctive cells. For each cell, a spike raster as a function of position (top), a spatial firing rate map (second row), the corresponding position firing-rate heat map (third row), and the unfolded view rate map (bottom) are shown. Text in red indicates the SI score for the position map and the VI score for the view map, each with its permutation-test p-value for the displayed condition. **(B)** Proportion of Conjunctive cells within each task condition (chi-square test; n.s., not significant). **(C)** Distributions of SI (position) and VI (view) scores for all Conjunctive cells, pooled across task conditions. **(D)** Distribution of position-firing stability for Conjunctive cells, quantified as Pearson correlation coefficients between position rate maps from odd and even laps; cells classified as stable (brown) or unstable (grey) based on a permutation test (p < 0.05). **(E)** Distribution of view-firing stability for Conjunctive cells, quantified as Pearson correlation coefficients between view rate maps from odd and even laps; cells classified as stable (green) or unstable (grey) based on a permutation test (p < 0.05).

We assessed the reliability of the two representations separately using odd-even lap correlations for position and view rate maps. Conjunctive cells were highly stable in both domains, with 88.6% showing stable position tuning (Fig. 7D) and 71.4% exhibiting stable view tuning (Fig. 7E). These findings identify a distinct hippocampal population in which information about self-location and visually sampled space is jointly and reliably represented.

### Population decoding separates position and view ensembles

Finally, the single-neuron dissociation between position and view coding was examined at the population level using Bayesian decoding. In each task condition, position could be decoded accurately from the recorded population, with decoded location closely tracking the animal’s actual location along the track (Fig. 8A). To test whether position-coding cells specifically supported this performance, decoding accuracy was compared across two ensembles: a position ensemble composed of Position and Conjunctive cells and a non-position ensemble composed of View and non-spatial cells. The position ensemble outperformed size-matched non-position ensembles in every condition, with error advantages of 11.17 (F-O; 95% CI [7.47, 14.52]), 7.01 (F-I; [4.96, 9.17]), 5.65 (C-O; [2.91, 8.94]), and 7.46 (C-I; [5.17, 9.77]) bins (Fig. 8B). Consistent with this, spike-train shuffling in the position ensemble strongly degraded decoding (red in Fig. 8C), whereas shuffling non-position cells had a much smaller effect (blue in Fig. 8C), indicating that accurate position decoding depended primarily on the activity of position-coding neurons.

**Figure 8.**
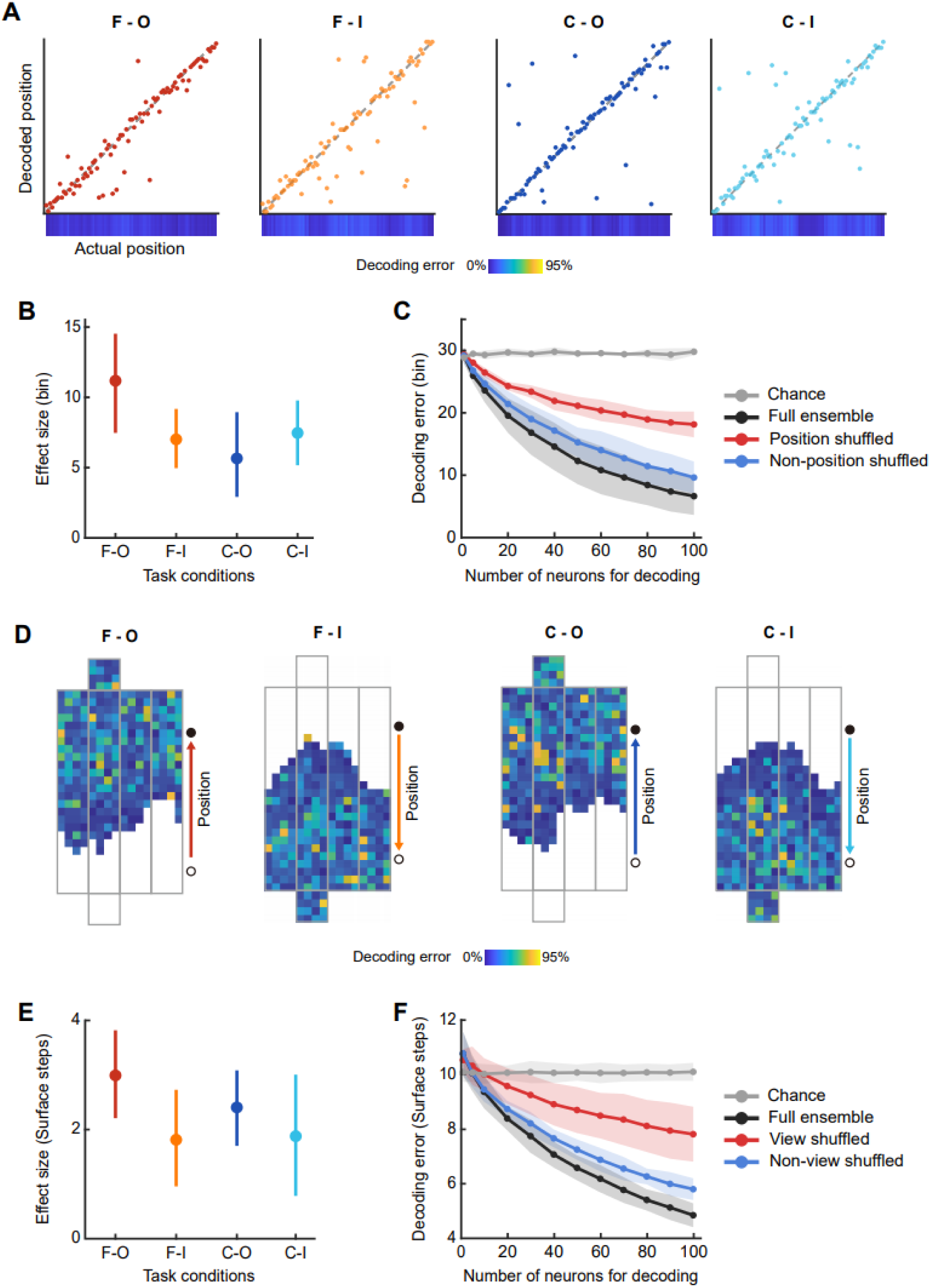
Functional contribution of position and view ensembles for population decoding. **(A)** Position decoding from the full recorded population. For each task condition, decoded position is plotted against actual position for held-out 250-ms samples from the Bayesian decoder, using all recorded cells in that condition. The diagonal line indicates no decoding error. The heat map below each scatter plot shows the mean decoding error at each position bin along the track. **(B)** Decoding advantage of the position ensemble (Position + Conjunctive cells). For each task condition, position-decoding error from the position ensemble is compared with that from a size-matched ensemble drawn from the non-position cells (View + Null). Filled circles indicate the mean difference in decoding error (comparison ensemble − position ensemble; *y*-axis, effect size in bins), and error bars indicate 95% confidence intervals across resampling repetitions; values > 0 indicate better (lower-error) decoding by the position ensemble. **(C)** Neuron-dropping analysis under selective spike-train shuffling. Grand-mean decoding error across the four task conditions is plotted as a function of the number of neurons. At each population size *n*, *n* cells were sampled from the full population and decoded under four manipulations: Full ensemble (black), all spike trains intact; Position shuffled (red), spike trains of the position ensemble shuffled, abolishing their tuning while retaining the cells in the decoded population; Non-position shuffled (blue), spike trains of the remaining non-position cells shuffled; Chance (grey), all spike trains shuffled. Shaded bands indicate 95% confidence intervals (*t*-distribution, *N* = 4 conditions). **(D)** View decoding from the full recorded population. For each task condition, the mean view-decoding error for each location on the unfolded view box is shown as a heat map. **(E)** Decoding advantage of the view ensemble (View + Conjunctive cells); View-decoding error from the view ensemble is compared with that from a size-matched ensemble drawn from non-view cells (Position + Null), with conventions and statistics as in (B) (units in surface steps). (F) Neuron-dropping analysis under selective spike-train shuffling for view decoding, with conventions as in (C). View shuffled (red), spike trains of the view ensemble shuffled; Non-view shuffled (blue), spike trains of the remaining non-view cells shuffled; units in surface steps.

An analogous result was obtained for spatial view. View could be decoded accurately across conditions from the full recorded population (Fig. 8D). A view ensemble composed of View and Conjunctive cells outperformed size-matched non-view ensembles in all conditions, with error advantages of 2.99 (F-O; 95% CI [2.61, 3.94]), 1.81 (F-I; [0.96, 2.73]), 2.41 (C-O; [1.70, 3.09]), and 1.88 (C-I; [0.78, 3.01]) surface steps (Fig. 8E). As in the position analysis, shuffling spikes in the view ensemble impaired decoding much more strongly than shuffling non-view cells (Fig. 8F). These population analyses demonstrate that the dissociation between position and view coding extends beyond single-cell classifications. Rather, the two forms of spatial information are carried preferentially by partially distinct ensembles that operate concurrently within the same hippocampal population.

## Discussion

The present findings show that the primate hippocampus contains coexisting but dissociable neural codes for self-location and spatial view during virtual navigation. Combining a field-of-view framework for gaze reconstruction with generalized linear modeling revealed three coexisting populations: Position cells with hallmark properties of conventional place cells, View cells forming stable representations of allocentric visual space, and a relatively small population of Conjunctive cells encoding both. Population decoding confirmed that this dissociation extends to the ensemble level, with position and view information carried preferentially by their respective ensembles. The fact that robust position codes coexist with spatial view codes during visually enriched, task-demanding virtual navigation suggests that the view-dominant description of primate hippocampal spatial coding may reflect, at least in part, methodological and task factors rather than an intrinsic constraint of the system.

One central implication is that spatial coding in the primate hippocampus cannot be reduced to a single dominant reference frame. Earlier primate studies either reported location-specific firing without fully dissociating it from gaze^35–37,40–42^, or emphasized view-based or mixed selectivity^11,12,15,16^. Our data extend these accounts by showing that a substantial population of neurons is better explained by position even after view-dependent sampling is explicitly modeled. During movement along a linear track, an animal’s location and the portion of space it views are intrinsically coupled, so position-related firing can be mistaken for view coding and vice versa; separating the two requires a model that estimates each contribution while controlling for the other. The current modeling framework addressed this problem by forcing position, view, and conjunctive explanations to compete against one another within each task condition. Under this approach, a distinct population remained best explained by position alone, indicating that view-independent location coding can be recovered in the primate hippocampus when the relevant variables are explicitly dissociated. The primate hippocampus therefore appeared to maintain parallel representations of where the animal is and what part of the environment it is sampling visually.

The properties of Position cells further indicate that this code shared several key properties with classical place cells. First, positional firing was spatially localized and reliable within a session, and the population collectively tiled the linear track, a population-level signature of a place-cell-like map^43,44^. Second, tuning was strongly directional, with most cells preferentially encoding a single travel direction, as reported in rodent studies^25–28,45^.

Third, a substantial fraction remapped between the Forest and City contexts, consistent with the hallmark global remapping across distinct contexts in rodent work^29–33^. Together, these features suggest that the primate position code closely resembles the canonical properties ascribed to place cells, pointing to an allocentric, map-like coding mode in the primate hippocampus that resembles the rodent counterpart in both form and function.

The results also help clarify why previous primate studies may have reported relatively few neurons with strong position-dependent tuning^11,12,15,16^. We suggest that two features of the present paradigm likely contributed to the detection of dissociable position signals. First, environmental richness: rodent studies showed that structured, enriched environments increase place-cell prevalence and precision^46–49^. Moreover, given primates’ reliance on high-acuity vision, our naturalistic VR contexts with distinct landmarks and boundaries may have promoted finer position tuning compared with the sparser environments of earlier primate studies^11,12^. Second, behavioral demands that required contextual memory during navigation: tasks demanding context discrimination enhance place-cell recruitment in rodents^50,51^. In the current task, context was not a background variable but a behaviorally relevant feature that had to be sampled repeatedly to guide object choice. Under such conditions, positional and contextual information may be more strongly engaged than during passive viewing or less structured navigation.

This stands in contrast to earlier primate work, where the relative scarcity of place cells has often been attributed to species-level differences, such as anatomical distinctions and divergent sampling strategies (e.g., whole-body movement in rodents versus head- and gaze-based sampling in primates) that were taken to favor view-over place-based coding^19–24,52^. Our identification of view-independent Position cells with place-cell-like properties, together with recent reports of location-selective primate activity^36,37^, challenges this view-centric account; the fact that the coding categories persist when position and spatial view are pitted against each other in a single model strengthens this point. The relative prominence of position or view coding may therefore depend less on species than on task structure and environmental richness.

This interpretation does not imply that view coding is secondary. On the contrary, View cells formed the largest of the three spatial classes, exhibited stable tuning, and showed robust contextual reorganization. The prominence of this code is consistent with the importance of active visual sampling in primate behavior and with longstanding proposals that primate hippocampal representations are shaped by gaze-dependent exploration of the environment. What the present data add is that visual and positional codes should not be treated as mutually exclusive alternatives. Instead, both appear to contribute to a heterogeneous representational architecture in the primate hippocampus. Consistent with this perspective, the heterogeneous coding we observed here may provide an efficient substrate for goal-directed behavior, with position, view, and conjunctive coding acting as dynamically weighted modes within a shared representational space that is reconfigured by the hippocampus according to sensory availability, behavioral goals, and environmental constraints.

Several limitations should be considered when interpreting the current findings. First, the task was constrained to linear-track navigation, which does not capture the full complexity of natural navigation. Second, environmental richness and task demands were manipulated together rather than independently, making it difficult to assign their relative contributions to the observed coding structure. Third, population decoding was performed on pseudo-populations assembled across sessions, which characterizes the information available in the recorded population but not the exact moment-to-moment readout of simultaneously active ensembles. Despite these limitations, the present study provides evidence that the primate hippocampus supports concurrent yet dissociable coding of position and spatial view during navigation. The contrast between place-like and view-like signals in primate work may therefore reflect not only species differences but also differences in behavioral design, environmental structure, and analytical resolution. More broadly, our findings support the view that primate hippocampal representations are heterogeneous and compositional, with overlapping neural populations carrying complementary information about spatial position and visually sampled space.

## Methods

### Subjects

Two adult male rhesus macaque monkeys were used in this study: monkey N (12 years old, 11 kg) and monkey Y (14 years old, 10 kg). The monkeys had ad libitum access to food, provided twice daily, while water access was restricted. All experimental procedures were approved by the Institutional Animal Care and Use Committee of Sungkyunkwan University (SKKUIACUC2022-03-62-1, SKKUIACUC2024-10-49-1, SKKUIACUC2024-12-02-1, SKKUIACUC2025-10-69-1).

### VR setup

We developed a virtual reality (VR) setup for monkey experiments. The setup consisted of a curved LCD monitor, a joystick, an eye-movement recording system (EyeLink 1000 Plus; SR Research), a movable experimental acrylic chair, a head-restraint system, and a reward delivery system (Fig. 1A). A curved LCD monitor (800 mm × 1,400 mm) was used to present the VR environment. To provide a realistic visual experience, the VR environment was constructed using a high-resolution game engine (Unreal Engine 4.14.3; Epic Games, Inc.).

The environment comprised two naturalistic spatial contexts, referred to as the Forest and City contexts, each containing both proximal and distal landmarks (Fig. 1A, B). The head-fixation system consisted of a frame mounted on the monkey’s head that was connected to a frame on the experimental chair. Bilateral fixation frames provided stable head restraint, ensuring that the monkeys’ head orientation was maintained facing the LCD monitor throughout the experiment. The monkeys performed navigation and behavioral responses using a joystick. Movement signals generated by joystick manipulation were transmitted to the computer via an Arduino Mega2560 and synchronized with the monkeys’ movements in the virtual reality environment using MATLAB R2021a (MathWorks) and Unreal Engine. Eye movements were simultaneously recorded using an infrared eye-tracking camera. The reward delivery system consisted of a licking tube mounted on the experimental chair and positioned in front of the monkey. Correct joystick responses during the contextual object choice trials (described in detail below) triggered a solenoid valve (VA212-3N; Aonetch) to dispense a 10 µl water reward by opening the valve for 0.1 s. Reward delivery was controlled by Unreal Engine via an Arduino Mega2560 interface.

### Shaping

After several weeks of handling, the monkeys were introduced to the VR apparatus for habituation and subsequently trained to navigate a virtual environment using a joystick. During this period, the Forest and City contexts were pseudo-randomly presented. The map comprised an inter-trial tunnel, a linear track, and a turning platform (‘rotary’). The monkeys were habituated to a stereotyped navigation sequence in which they initiated movement from the inter-trial tunnel, traversed the linear track, experienced a passive 180° rotation at the rotary, and returned to the inter-trial tunnel; this complete sequence was defined as a lap.

Following successful navigation shaping (100 laps within 120 minutes on two consecutive days), the monkeys were trained to perform a choice response task within the same map.

Upon reaching predefined locations during outbound and inbound navigation, a heart-shaped 3D object appeared on either the left or right side of the track. The monkeys obtained the reward by manipulating a joystick in the direction of the presented object. After completing 210 trials within 120 minutes on two consecutive days, the monkeys learned the final shaping of choosing one of the two objects based on their visual properties. Objects were presented with either full opacity (100%) or low opacity (10%), and the monkeys were required to select the fully opaque object. Monkeys that completed more than 210 trials within 120 minutes and achieved an accuracy exceeding 80% on two consecutive days advanced to the main task.

### Context-dependent object-choice task

In the main task, the monkeys navigated the same contexts described above and learned the associations between contexts and objects. The two contexts were presented in a pseudo-random order. Objects were divided into two categories: one object group (i.e., pumpkin, jellyfish, and pizza) was rewarded in the Forest context, whereas the other object group (i.e., donut, turtle, and octopus) was rewarded in the City context; objects were not rewarded in the opposite context (Fig. 1B). When the monkeys reached predefined trial locations during navigation, a contextual object-choice trial was initiated (Fig. 1C, Supplementary Fig. 2A).

At trial onset, locomotion was halted and a red circle was presented at the center of the screen as a ready cue. Following a variable delay of 300–600 ms after ready cue onset, two objects—each randomly selected from one of the two object categories—were presented on the left and right sides of the ready cue. After an additional delay of 300–600 ms, the ready-cue changed from a red circle to a blue circle, signaling go-cue onset. Following go-cue onset, the monkeys were required to select the object rewarded in the current context by manipulating the joystick left or right. Correct choices were followed by an auditory feedback tone (1300 Hz) and delivery of a water reward (solenoid valve opened for 0.1 s; 10 µl) through the reward tube. Incorrect choices resulted in a low-frequency auditory feedback tone (140 Hz) without reward. Within a single lap, eight trials were conducted, with four trials occurring during outbound and the other four during inbound navigation.

Training progressed in a stepwise manner with increasing object-set complexity. The monkeys were first trained with a single object pair (pumpkin and donut), followed by a second object pair (jellyfish and turtle). Training then proceeded with four objects combining the first and second object pairs, followed by a third object pair, and finally with all six objects. Advancement to the next training stage required completion of at least 100 laps (800 trials) with more than 80% accuracy on two consecutive days at each stage.

For all behavioral and neuronal analyses, we included only sessions in which the monkeys achieved more than 80% correctness for each object and completed at least 72 laps. This criterion ensured that all possible combinations of two contexts, four trial locations, two navigation directions (outbound/inbound), six object identities, and two object locations (left/right) were presented, allowing each condition to be experienced at least twice.

### Surgery

Prior to surgery, implantation coordinates for the head post and recording chamber were calculated based on the location of the hippocampus, which was identified using 3T and 7T MRI scans. Surgeries were performed under isoflurane anesthesia in accordance with each monkey’s individual training schedule. For monkey N, the first surgery involved implantation of a custom-made titanium head post for head fixation. Ten titanium bone screws were inserted into the skull and cemented together with the head post to form a stable base supporting the implant. This procedure mechanically integrated the head post, skull, and bone screws, resulting in a rigid and stable head-fixation system. The second surgery involved implantation of a recording chamber. A custom-built recording chamber (23 mm × 25 mm; Ultem) was affixed to the skull and secured using seven ceramic screws and dental cement.

The chamber was designed to accommodate a recording grid (19 mm × 21 mm). When experiments were not being conducted, a chamber cover was secured to the top of the chamber. For monkey Y, implantation of the head post and recording chamber was performed during a single surgical session using the same specifications and procedures as monkey N.

### Electrophysiological recording

After sufficient postoperative recovery, MRI scans were acquired to determine the depth and coordinates from the recording chamber to the hippocampus. A recording grid structure containing a contrast agent was placed within the recording chamber for each recording session and was installed and removed at every session; the recording chamber was thoroughly cleaned according to established protocols before and after each session to prevent infection. The recording grid consisted of a rigid guide-tube holder with regularly spaced holes arranged in a 19 × 21 array, designed to precisely guide electrode trajectories toward the target region. Each hole in the grid served as a fixed coordinate, allowing reproducible and stable electrode insertions across recording sessions.

For each recording session, hippocampal target coordinates were selected based on MRI images. Electrodes were mounted on a motorized electrode manipulator pencil (MEM pencil, Thomas RECORDING GmbH) equipped with a guide tube. Prior to electrode insertion, the electrodes and guide tubes were sterilized by immersion in alcohol for at least 30 minutes. The recording grid was then installed in the recording chamber, and the sterilized electrode was inserted through the recording grid. Electrode advancement and retraction were controlled via a microprocessor motor control unit (MCU-2, Thomas RECORDING GmbH; speed of 15 µm/s). After reaching the target depth, a waiting period of at least 30 minutes was allowed to permit recovery from transient brain compression. Subsequently, cell hunting was performed by slowly advancing the electrode at a rate of 2–5 µm/s. Once a well-isolated neuronal signal was detected, recordings commenced after an additional stabilization period of at least 15 minutes.

Neural activity was recorded using a Digital Lynx acquisition system (Neuralynx). Signals were amplified 1,000–10,000-fold and band-pass filtered between 300 and 6,000 Hz for spike detection. Spike waveforms exceeding a manually adjusted threshold (typically 30–100 µV) were digitized at 32 kHz and time-stamped.

At the end of each recording session, the electrode was retracted at a speed of 15 µm/s. The electrode and recording grid were then removed, the recording chamber was thoroughly cleaned according to established protocols, and the chamber cover was reinstalled to seal the recording chamber.

All data were collected after completion of behavioral training (i.e., completed more than 100 laps within 60 minutes and achieved over 80% correctness for each object across two consecutive days). Neuronal and behavioral data—including trial events, the monkey’s position within the virtual environment, and joystick manipulation—were recorded in synchrony with the VR system at 30 Hz. Neural recordings were obtained using either a single electrode or a tetrode (MEM pencil, Thomas RECORDING GmbH).

#### Unit isolation

All single units were manually isolated using different software depending on electrode type: Offline Sorter (Plexon) for single electrodes and a custom program (WinClust) for tetrodes. Peak amplitude was used as the primary criterion for spike sorting, along with additional waveform features including energy and peak-to-trough latency. Units were excluded if more than 1% of spikes occurred within the refractory period (1 ms).

#### Unit filtering

Only units from sessions with more than 72 valid laps were included, after excluding laps invalidated by experimenter intervention or mechanical issues. A total of 253 units were recorded from these valid sessions. Based on cluster isolation, 156 units were classified as single units and 97 units as multi-unit activity. Units were assigned to multi-unit activity when cluster boundaries were poorly separated or when multiple distinct waveform shapes were present within a single cluster. Among the 156 single units, 12 were identified as fast-spiking neurons, defined by a mean firing rate greater than 10 Hz and a spike width shorter than 350 µs, and were excluded from further analyses. During recordings, we intentionally targeted putative principal cells and avoided fast-spiking neurons based on established electrophysiological characteristics (e.g., firing rate and spike waveform), to preferentially sample excitatory neurons.

For putative complex-spiking neurons, all analyses were restricted to spikes and behavioral data recorded during active navigation. Periods with movement speeds lower than 15 cm/s in the VR environment, epochs corresponding to contextual object-choice trials, and positions outside the linear track (700–9,500 cm) within each context were excluded.

Neurons that emitted fewer than 100 spikes during the included periods across all laps were also excluded. After applying these criteria, 120 neurons were retained for analyses (Supplementary Table 1). For condition-wise analyses, neurons were further required to fire more than 30 spikes within a given task condition; cells not meeting this criterion in a condition were excluded from that condition’s analyses. A total of 113 neurons exceeded this threshold in at least one of the four conditions and were included in the condition-wise analyses (F-O, n = 102; F-I, n = 104; C-O, n = 104; C-I, n = 103).

#### Preprocessing of eye-movement data

Raw eye-movement data along the horizontal (X) and vertical (Y) axes were first preprocessed to reduce high-frequency noise. Specifically, the signals were low-pass filtered using a second-order Butterworth filter with a cutoff frequency of 25 Hz, applied in a zero-phase (two-pass) manner to avoid phase distortion. The sampling rate of the eye-position data was 1,000 Hz. Filtered eye-movement data were converted from linear displacement to angular units (degrees of visual angle) using the arctangent of the scaled eye-position displacement relative to the eye-to-monitor distance. Instantaneous eye velocity was computed as the first temporal derivative of angular eye position for horizontal and vertical components and expressed in degrees per second. Eye acceleration was calculated analogously as the first temporal derivative of eye velocity, yielding units of degrees per second squared. Horizontal and vertical components of velocity and acceleration were combined to form two-dimensional vectors.

Saccades were detected using a multi-step algorithm based on eye acceleration, velocity, and movement direction. Candidate saccade periods were first identified by thresholding horizontal and vertical eye acceleration signals. Samples exceeding an acceleration threshold of 1000 deg/s² in either axis were marked as saccade candidates. Short gaps within candidate segments shorter than 40 ms were merged, and candidate segments separated by inter-saccadic intervals shorter than 10 ms were combined to form preliminary saccade epochs. For each candidate saccade, two-dimensional eye velocity was computed, and the time point of peak velocity was identified. The main saccade direction was estimated as the average of the sample-to-sample velocity direction at the peak velocity sample and its immediate neighboring samples. Saccade onset and offset were then refined using directional and velocity-based criteria. Boundaries were adjusted when the deviation from the main saccade direction exceeded 60°, or when deviations larger than 20° occurred for three consecutive samples. Additional refinement was applied based on sample-to-sample changes in movement direction using the same angular thresholds. Finally, saccade onset and offset were constrained by velocity criteria: saccade boundaries were defined as the points at which eye velocity fell below an absolute threshold of 30 deg/s, or below 20% of the peak velocity for that saccade. Samples between the refined onset and offset were labeled as saccades, yielding a binary saccade index for each time point. Eye-movement samples classified as saccadic movements were excluded from subsequent analyses. All remaining samples, including fixation and smooth eye-movement periods, were grouped as non-saccadic fixations and used for all eye-movement analyses reported in this study.

#### Constructing 3D gaze map

To reliably measure gaze positions within the 3D VR space, we introduced a virtual “view box” (length 13,000 cm, width 2,000 cm, height 2,000 cm; Fig. 1E) within each VR context. 2D eye positions recorded from the monkeys were transformed and projected onto the 3D coordinate of the view box. This projection was performed by computing the intersection between the virtual view box and rays defined by viewing angles derived from the monkey’s eye position in 2D coordinates, enabling the reconstruction of gaze positions within the 3D VR space.

To define the spatial extent of visual perception, the reconstructed gaze point was subsequently expanded into a field-of-view (FOV). Prior to FOV estimation, the entire view box was discretized into uniform three-dimensional spatial bins with a resolution of 500 cm. A viewing angle of ±60°, encompassing the monkey’s peripheral visual range, was applied to construct a 2D FOV boundary centered on the fixation point. This boundary was projected into the 3D space, and each spatial bin was classified according to whether it fell within the resulting FOV boundary based on geometric intersection criteria.

To further approximate the non-uniform nature of visual information processing across the visual field, a Gaussian weighting function (σ = 8) was applied with respect to the fixation point. For each spatial bin within the FOV boundary, the Euclidean distance from the fixation point was calculated, and a corresponding Gaussian value was assigned, such that bins closer to the fixation point were weighed more strongly while bins toward the periphery received progressively lower weights. This procedure yielded a three-dimensional, spatially weighted FOV representation that captured both the geometric extent of gaze and the graded decline of visual sensitivity across peripheral vision (Fig. 1F).

#### Decoding contexts with eye movements

To compare gaze behaviors across contexts, we performed a decoding analysis using gaze data as features, separately for each monkey. Within each session, we added fixation counts (448 spatial bins) from both outbound and inbound conditions for the Forest and City contexts separately. These counts were z-score normalized within each session to provide the feature vectors for subsequent analysis. We used multidimensional scaling (MDS) to visualize the clustering of Forest and City contexts based on gaze behaviors in three-dimensional space (Supplementary Fig. 4C). We then applied a linear SVM to perform binary classification of whether each gaze data is from Forest or City context (Supplementary Fig. 4D). We used 5-fold cross-validation to train and test the model. To test the significance of decoding results, we shuffled the context labels and performed the decoding 1000 times to construct surrogate distribution of the decoding results. p-values were determined by comparing with the surrogate distribution.

#### GLM-based cell classification

To dissociate position and spatial view at the single-neuron level, we fit Poisson GLMs to each neuron’s binned spike train, separately for each of the four conditions, comparing three encoding models: a position model (location parameterized by Gaussian basis functions along the track), a view model (gaze location parameterized as the bins of the unfolded view box), and a conjunctive model combining both.

Cross-validation was lap-based to avoid within-lap leakage: laps were split into 9 outer folds, and a nested 3-fold cross-validation on the training laps selected the ridge-penalty strength λ (from a logarithmically spaced grid) by minimizing held-out Poisson deviance. Sparsely sampled view bins (low cumulative occupancy) were masked to prevent unstable fits. For each test fold, encoding performance was quantified as the cross-validated log-likelihood gain per bin over a mean-rate null model (ΔLLH/bin).

Each neuron was classified by forward selection on the fold-wise ΔLLH/bin. The better single-variable model (position or view) was first tested against the null (one-tailed Wilcoxon signed-rank across folds; α = 0.05); neurons not exceeding the null were classified as non-spatial (Null). Otherwise, the conjunctive model was tested against the best single model and the neuron classified as Conjunctive if it significantly improved on it, or as a Position or View cell otherwise. Classification was performed independently in each condition.

Each model was also refit on all of a condition’s data for the rate map reconstructions shown in the figures; the significance of each fit was assessed by circularly shifting the spike train relative to the predictors (1,000 shuffles) and comparing the observed log-likelihood to the shuffle distribution.

#### Spatial information score (SI)

The linear track was divided into 89 spatial bins of 100 cm (700–9,500 cm) each, and the firing rate for each bin was calculated by dividing the spikes count occurring within the bin by the occupancy of that bin, yielding a spatial firing-rate map. After map construction, smoothing was applied using the Skaggs adaptive binning method. The spatial information of the single cell was calculated from the smoothed spatial firing-rate map according to the equation below^39^:

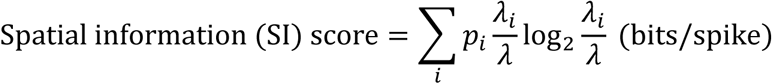

where i is the spatial bin index, p_i the occupancy probability of bin i, λ_i the mean firing rate in bin i, and λ the overall mean firing rate across bins. SI was computed separately for each of the four conditions (two contexts × two navigation directions). The significance of each SI score was assessed against a null distribution generated by shuffling spike times (n = 1,000 shuffles).

#### View information score (VI)

The firing rate of the spatial view-rate map was computed by dividing the Gaussian-weighted spike count of each bin of the view box by its corresponding Gaussian-weighted occupancy. The resulting map was smoothed using the Skaggs adaptive binning method. The spatial view information of each single cell was then calculated from the smoothed rate map for spatial view according to the equation below^39^:

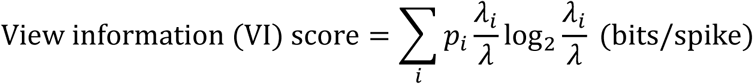

where *i* is the view box bin index, *p_i* the occupancy probability of bin *i*, *λ_i* the mean firing rate in bin *i*, and *λ* the overall mean firing rate across bins. VI was computed separately for each of the four conditions. As for SI, the significance of each VI score was assessed against a spike-shuffled null distribution (*n* = 1,000 shuffles).

#### Rate map stability and remapping

To assess the reliability of position and view tuning, we computed split-half (odd–even) stability. For each cell and condition, rate maps were constructed separately from the odd and even laps, and the stability index was the pixel-wise correlation between the two maps, taken over bins occupied in both halves (1D position maps or 2D view maps, as defined above).

Significance was assessed against a null distribution of 1,000 shuffles, each generated by circularly shifting the spike train by a random offset (10–90% of the session), then re-splitting the laps and recomputing the odd–even correlation. The *p*-value was the fraction of shuffles whose correlation met or exceeded the observed value; cells with p < 0.05 were classified as stable, and the remainder as unstable (Figs. 3E, 5E, 7D, E).

To quantify remapping, we compared a cell’s rate maps across the two directions (within a context; directional remapping) or across the two contexts (within a direction; contextual remapping), restricted to cells classified as a Position (or View) cell in both members of the pair. The across-direction or across-context correlation was tested against a null distribution generated by the same shuffling procedure (1,000 circular shifts); cells whose correlation was significant (p < 0.05) were classified as stable, and the remainder as remapping (Figs. 4C, F; 6C).

#### Bayesian decoding of position and spatial view

To quantify the spatial information encoded by the hippocampal population, we decoded position and spatial view from a pseudo-population of neural activity using a Bayesian maximum a posteriori (MAP) approach within 250-ms observation windows. For each condition, decoding used the per-condition populations defined in *Cell filtering*. For each window, synthetic spike counts were generated by sampling from a Poisson distribution, using each neuron’s rate map to model its firing probability at each spatial state (s). Assuming independent firing across neurons and a uniform prior P(s), the monkey’s spatial state was estimated from the likelihood P(n|s):

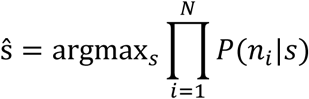

where ŝ is the decoded state (position or spatial view), n is the vector of simulated firing counts for N neurons, and P(n_i|s) is the Poisson probability of count n_i given state s. Decoding error was quantified as the distance between the actual and decoded state (position, track bins; spatial view, surface steps on the view box), and its significance was assessed against a surrogate distribution generated by shuffling firing rates (n = 1,000).

To test the specific contribution of each spatial code, we compared decoding from the functional ensembles with size-matched comparison ensembles. For position, the position ensemble (Position and Conjunctive cells) was compared with an equally sized ensemble drawn at random from the remaining, non-position (View and Null) cells; for view, the view ensemble (View and Conjunctive cells) was compared with a size-matched ensemble of non-view (Position and Null) cells. Across many resampling repetitions, the effect size was defined as the mean difference in decoding error (comparison ensemble − functional ensemble), with 95% confidence intervals across repetitions; positive values indicate lower-error decoding by the functional ensemble (Fig. 8B, E).

Finally, a neuron-dropping analysis was performed under selective spike-train shuffling. At each population size, neurons were subsampled from the full population (100 repetitions) and decoded under four manipulations: all spike trains intact (Full ensemble); the spike trains of the position (or view) ensemble shuffled, abolishing their tuning while retaining the cells in the decoded population (Position/View shuffled); the spike trains of the remaining non-position (non-view) cells shuffled (Non-position/Non-view shuffled); and all spike trains shuffled (Chance). This quantified how decoding error scaled with the number of neurons and which ensemble drove the decoding (Fig. 8C, F).

## Acknowledgments

This research was supported by the National Research Foundation of Korea 2018R1A4A1025616, 2019R1A2C2088799, 2021R1A4A2001803, 2022M3E5E8017723, RS-2024-00452391, RS-2025-02303740 and BK21 FOUR program. This work was supported by Mid-Career Bridging Program through Seoul National University and Institute for Basic Science IBS-R015-D1 and IBS-R015-D2.

