## Supplemental data for "Coexisting but dissociable place and spatial view codes in the primate hippocampus"

**Functional dissociation of place and spatial view codes in primate hippocampus**

^†^**Corresponding author:**

Inah Lee

Joonyeol Lee

**Supplementary information**

| **Number of recorded cells** | | | |
| --- | --- | --- | --- |
| **Single-unit** | | | **Multi-unit** |
| 156 | | | 97 |
| **Complex spiking neurons** | | **Fast spiking neurons** |  |
| 144 | | 12 |  |
| **Number of spikes > 100** | **Number of spikes ≤ 100** |  |  |
| 120 | 24 |  |  |

**Supplementary Table 1. Number of recorded units at each filtering step.** Units obtained from two monkeys were classified as single units or multi-unit activity following cluster isolation criterion. Single units were further categorized as complex-spiking or fast-spiking neurons based on spike width and mean firing rate criteria. Complex spiking neurons with insufficient spike counts were also excluded from subsequent analyses to ensure reliable estimation of spatial tuning (see Methods for details).

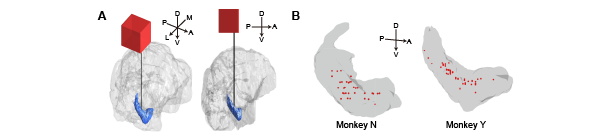

**Supplementary Figure 1. Reconstruction of recording sites in the primate hippocampus. (A)** 3D reconstruction of the macaque brain derived from structural MRI scans. The right hippocampus is highlighted in blue. The red square box indicates the recording chamber placement, and the black vertical line represents a possible electrode trajectory. D, dorsal; V, ventral; A, anterior; P, posterior; M, medial; L, lateral. **(B)** Summary of estimated recording sites for Monkey N (left) and Monkey Y (right) projected onto the reconstructed hippocampus.

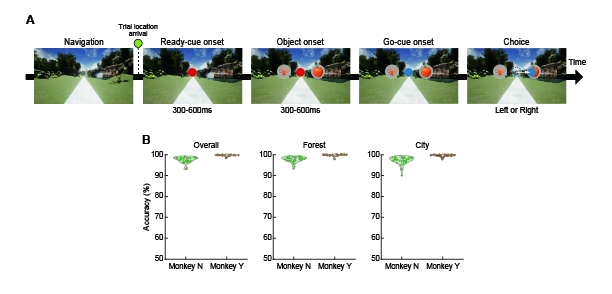

**Supplementary Figure 2. Contextual memory task during virtual navigation. (A)** Schematic of a single trial, triggered when the monkey arrived at a predefined trial location. Following a go-cue, the monkey manipulated a joystick (left or right) to select one of two objects associated with the current context. **(B)** Behavioral performance in the context-dependent object-choice task. Session-averaged accuracy is shown for each monkey. Both animals maintained performance above 90% in both contexts.

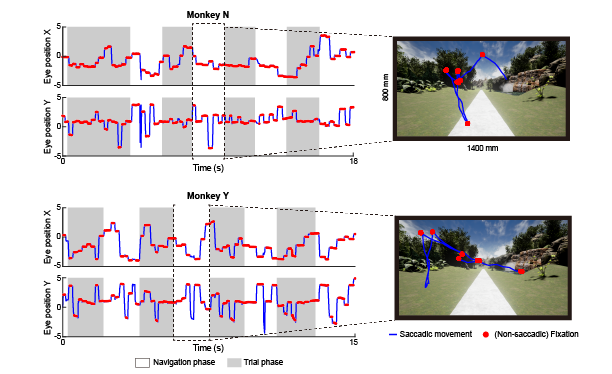

**Supplementary Figure 3. Classification of saccadic movements and fixations.** Spatiotemporal traces of horizontal (X) and vertical (Y) eye positions (left). Eye movements during the navigation phase (white) were only considered for the subsequent analyses. Representative eye movements projected onto the LED TV (right). Only non-saccadic fixations were used for the 3D reconstruction of gaze behavior in Fig. 1.

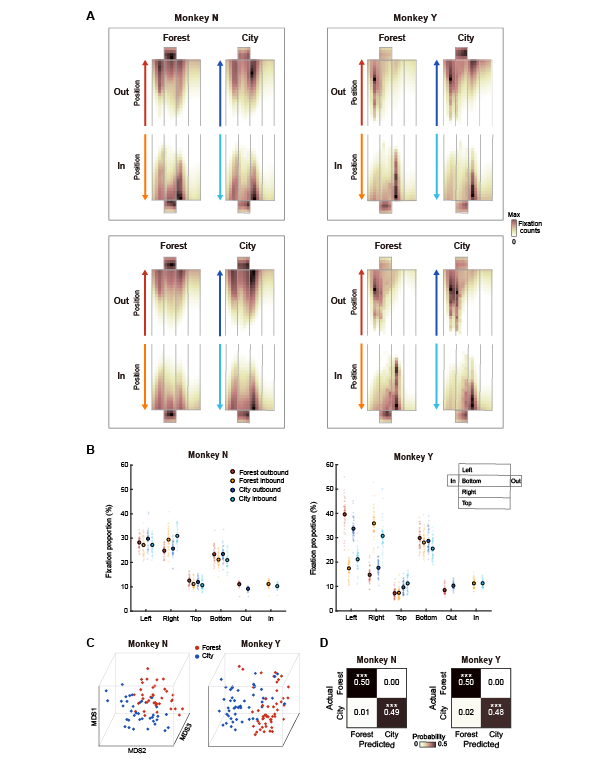

**Supplementary Figure 4. Context-dependent gaze patterns during virtual navigation. (A)** Representative unfolded 2D gaze maps illustrating distinct gaze patterns across contexts (Forest and City). Maps from single representative sessions of Monkey N and Monkey Y are organized by context (Forest, City) and travel direction (Outbound, Inbound). **(B)** Quantification of fixation proportions across the six surfaces of the virtual view box. Each small dot represents the fixation proportion from a single session, and large circles indicate the average across all sessions. (**C**) Low-dimensional projections of gaze patterns obtained using multidimensional scaling (MDS) to visualize clustering of gaze behavior in the Forest and City contexts. Each dot represents the gaze pattern of a single session. (**D**) Performance of a linear support vector machine decoding context (Forest vs. City) from gaze maps. Confusion matrices show significantly high classification accuracy for both monkeys. ***p < 0.001.
